# High-throughput screening of functional neo-antigens and their specific TCRs via the Jurkat reporter system combined with droplet microfluidics

**DOI:** 10.1101/2023.02.20.529171

**Authors:** Yijian Li, Jingyu Qi, Yang Liu, Yuyu Zheng, Haibin Zhu, Yupeng Zang, Xiangyu Guan, Sichong Xie, Hongyan Zhao, Yunyun Fu, Haitao Xiang, Weicong Zhang, Huanyi Chen, Huan Liu, Yuntong Zhao, Yu Feng, Fanyu Bu, Yanling Liang, Yang Li, Qumiao Xu, Ying He, Li Sun, Longqi Liu, Ying Gu, Xun Xu, Yong Hou, Xuan Dong, Ya Liu

## Abstract

T-cell receptor (TCR)-engineered T cells can precisely recognize a broad repertoire of targets derived from both intracellular and surface proteins of tumor cells. TCR-T adoptive cell therapy has shown safety and promising efficacy in solid tumor immunotherapy. However, antigen-specific functional TCR screening is time-consuming and expensive, which limits its application clinically. Here, we developed a novel integrated antigen-TCR screening platform based on droplet microfluidics technology, enabling high-throughput peptide-major histocompatibility complex (pMHC) library-to-TCR library screening with high sensitivity and low background signal. We introduced DNA barcoding technology to label peptide antigen candidate-loaded antigen-presenting cells (APCs) and Jurkat reporter cells to check the specificity of pMHC-TCR candidates. Coupled with the next-generation sequencing pipeline, interpretation of the DNA barcodes and the gene expression level of the Jurkat T-cell activation pathway provided a clear peptide-MHC-TCR recognition relationship. Our proof-of-principle study demonstrates that the platform could achieve unbiased pMHC-TCR library-on-library screening, which is expected to be used in the cross-reactivity and off-target testing of candidate pMHC-TCR libraries in clinical applications.

## Introduction

T-cell immunotherapy is one of the most promising cancer immunotherapies^1^, which includes tumor-infiltrating lymphocyte (TIL) therapy^2–5^, chimeric antigen receptor T-cell (CAR-T) therapy^6–9^, and T-cell-receptor T-cell (TCR-T) therapy^10–14^, and has been shown to achieve tumor regression. For solid tumors, TCR-T appears to be the most promising technology in the field of solid tumor targeted therapy^10,12^. TCR-T cells can recognize a wide range of tumor neo-antigens, not only those on the cell surface but also intracellular neo-antigens presented by the major histocompatibility complex (MHC)^15–17^. When a TCR binds to its cognate peptide–MHC complex (pMHC), the associated CD247 (CD3?) chains dimerize to initiate downstream signaling, leading to rapid gene expression driven by the transcription factors NF-κB, AP-1, and NFAT^18^. The affinity of the TCR can also be modified, allowing the optimal balance between the potency and off-target effects to be found^19^. Moreover, TCR-T cells may exhibit better tumor infiltration because of this fine-tuned affinity and would not tend to remain at the tumor periphery, making these cells more efficient for treating solid tumors^20,21^.

The conventional approaches for identifying functional antigen-specific T-cell clones, however, usually take several months, and are resource-intensive^22^ and expensive^23^, potentially leading to the optimal treatment time for cancer patients being missed. Billions of different TCR clonotypes are in humans, but only a few are specific for certain antigens^24,25^. Minimizing the risk of cross-reactivity and off-target recognition is also key for clinical application. These factors may have significant impacts on the treatment of cancer patients. There is a need for a rapid high-throughput system for screening functional TCRs and paired antigens, as well as the cross-reactivity of TCRs, which could significantly decrease the waiting time for cancer therapy.

Some high-throughput methods for T-cell antigen identification have been established. TCR and pMHC affinity-based assays, such as pMHC multimers^26,27^, yeast display^28,29^, TCR-pMHC interaction Koff-Rate^30^, YAMTAD^31^, and dextramer-TCR binding assays^32^, help to screen TCR–pMHC interactions by evaluating the binding affinity between TCR and pMHC molecules. However, these approaches are associated with the challenges of low efficiency in pMHC multimer production and instability of pMHC multimers^33^. Moreover, these assays can identify high-affinity TCRs, which would significantly increase the risk of cross-reactivity and off-target recognition^34,35^. Although Pei et al.^36^ recently reported that dorimer (DNA origami-based pMHC multimer) was capable of binding low-affinity TCR, the throughput of screening was still limited. Finally, affinity might not be a reliable indicator of T-cell response^37,38^ because TCR–pMHC interaction may be related to high binding affinity but not specific stimulatory interaction. It may thus be more accurate to identify the TCR–pMHC interaction based on the biological phenomenon of cell–cell interaction^37,38^.

Assays based on T-cell function or TCR active signaling pathway, such as trygocytosis^39,40^, T-scan^41^, “catch bound” ^42^, cytokine-capturing^43^, or combining single-cell droplet microfluidics with TCR active signaling pathway^44,45^, can be used to screen functional TCRs or antigens. These high-throughput screening methods have made a major contribution to the progress of antigen or TCR recognition, but they still face the challenge of limited throughput. They cannot be used to perform unbiased screening of a functional TCR and antigen simultaneously when candidate TCR libraries and peptide libraries are available; evaluating the interaction between two such libraries would thus be time-consuming^21^. They can also not achieve high-throughput screening of cross-reactivity and off-target recognition between candidate antigens and TCRs in two libraries. Therefore, simultaneously identifying the relationship between antigens and their specific TCRs via high-throughput screening is still difficult, while major obstacles to clinical application include low specificity of immune function and off-target cytotoxicity.

As an emerging technology, microfluidic technology is favored by researchers due to its miniaturization, integration, high sensitivity, and small sample consumption. Owing to its great potential in numerous fields, it has developed into an interdisciplinary field integrating biology, medicine, chemistry, electronics, materials science, and other fields^46^. Segaliny et al.^44^ performed functional screening of TCR-T cells using droplet microfluidics and monitored the activation of individual TCR-T cells in real time after the identification of target tumor cells. Downstream molecular analysis was performed using single-cell RT-PCR and TCR sequencing to confirm specificity. Although this research enabled observation of the activation of TCR-T cells in real time, the subsequent single-cell recovery steps were complicated and had low throughput. Similarly, Wang et al.^45^ used droplet microfluidic technology to co-encapsulate antigen-presenting cells (APCs) and TCR-T cells in droplets to allow stimulation at the single-cell level. Functional TCR-T cells were subjected to high-throughput screening by di-electrophoresis in microfluidic channels. The strategy of combining scRNA-seq would help to reveal the molecular mechanism of TCR–pMHC recognition. However, the pairing relationship between TCR-T cells and APCs could not be identified and traced back in a high-throughput identification of the TCR–pMHC interaction. Therefore, the introduction of microfluidic technology may help us achieve high-throughput screening of functional neo-antigens and their specific TCRs, and accelerate research on TCR-T therapies for treating solid tumors.

Here, we developed a high-throughput platform, which combines a Jurkat reporter system^47–52^, DNA barcoding^53^, and droplet microfluidics^45^, to enable the library-on-library screening of functional antigens and their specific TCRs. Jurkat reporter cells expressing candidate TCRs and antigen-presenting cells (APCs) pulsed with different antigens were labeled with different barcodes through Lipofectamine 3000 transfection. By droplet microfluidics, a Jurkat reporter TCR cell and APC were encapsulated in the same droplet, into which lysis buffer was injected to release the DNA barcode and mRNA without breaking the cell pairing in the droplet. DNA barcodes and transcription levels of genes involved in the T-cell activation pathway were then analyzed by single-cell RNA-sequencing technology, to interpret the cell pairing in each droplet and the specific recognition relationship between the antigen and its functional TCR. Our proof-of-principle study demonstrates that this high-throughput platform can be used for unbiased library-on-library screening of antigens and their functional TCRs simultaneously (two peptides and two TCRs) with high sensitivity and specificity. The high-throughput screening platform provides a powerful tool for simultaneously identifying tumor antigens and specific TCRs, which could significantly decrease the time required for screening functional antigens and specific TCRs, limiting the delay before cancer treatment can be initiated. This platform could also be used to screen the cross-reactivity and off-target recognition of candidate antigens and TCRs in libraries before clinical application to promote tumor immunotherapy.

## Results

### The workflow of high-throughput screening of functional neo-antigens and their specific TCR platform

To develop the high-throughput screening platform, we combined a Jurkat reporter system^47–52^, DNA barcoding^53^, and droplet microfluidics^45^ to enable the library-on-library screening of functional antigens and their specific TCRs. First, these candidate TCRs obtained from tumor-infiltrating lymphocytes (TILs) were expressed in Jurkat reporter cells (Figure 1A). Second, through Lipofectamine 3000 transfection, the APCs pulsed with different antigens and the Jurkat reporter TCR cells were labeled with different barcodes (Figure 1B). Third, a Jurkat reporter TCR cell and APC were encapsulated in the same droplet by droplet microfluidics and then incubated for 4 h (Figure 1C). Fourth, lysis buffer was injected into these droplet cells to lyse them, after which the DNA barcode and mRNA were released and captured by RNA beads without breaking the cell pairings in the droplets (Figure 1D). The capture tail sequence in the DNA barcodes and the poly-A sequence in the mRNAs enabled their capture by RNA beads, followed by reverse transcription together during the scRNA-seq process. Fifth, analysis of DNA barcodes allowed determination of the cell pairing in each microfluidic droplet. Finally, the transcription levels of genes involved in the T-cell activation pathway were analyzed through single-cell RNA-sequencing technology to interpret the specific recognition relationship between the antigen and its functional TCR (Figure 1E). The high-throughput platform can be used for simultaneous, unbiased, library-on-library screening of an antigen and its functional TCR with high sensitivity and specificity. This platform can significantly decrease the time required for screening functional antigens and specific TCRs, but it could also be used to screen the cross-reactivity and off-target recognition of candidate antigens and TCRs in libraries before clinical application to promote tumor immunotherapy.

**Figure 1.**
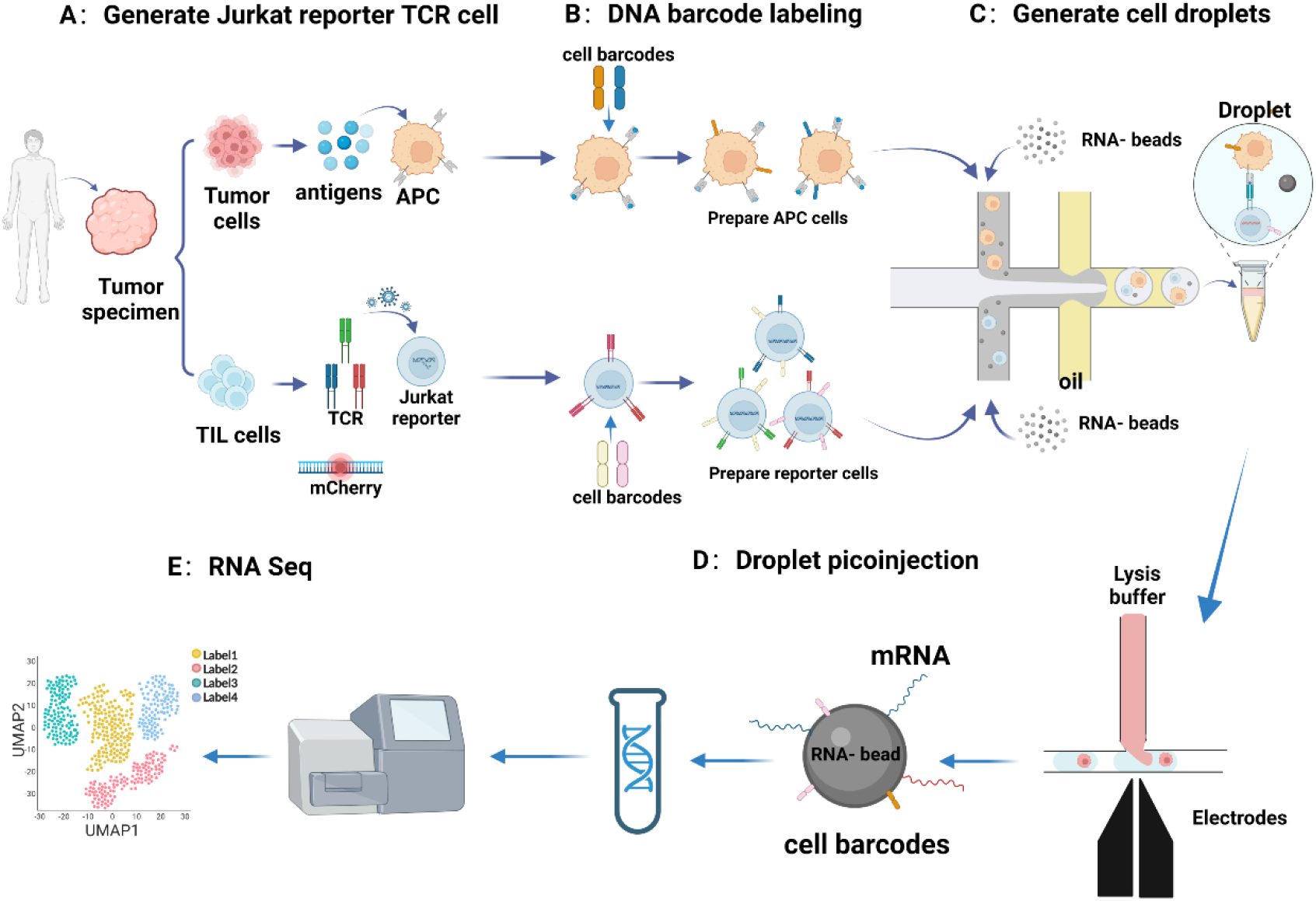
Schematic diagram of high-throughput screening of functional TCRs. Step 1: The candidate TCR was overexpressed by lentivirus on Jurkat reporter cells in which endogenous MHC-I and TCR had been knocked out through CRISPR/Cas9. The candidate peptide was applied in a pulsed manner to APC cells, in which the corresponding MHC-I molecule was overexpressed by lentivirus. Step 2: The cells were labeled with DNA barcode through Lipofectamine 3000 transfection. Step 3: Droplet generation, and co-encapsulation of cells and RNA beads. Step 4: Droplet picoinjection, involving injection of lysis buffer into these droplets for scRNA-seq. Step 5: Single-cell RNA-sequencing, transcript capture, library preparation, and scRNA-seq to analyze the functional antigens and specific TCRs. Validation of function of antigen and TCR using primary CD8^+^ T cells.

### Droplet generation and picoinjection

The droplets containing RNA beads and cells were generated by applying negative pressure to the droplet-generating chip, as previously described (Combinatorial perturbation sequencing on single cells using microwell-based droplet random pairing). As shown on the left of Figure 2A, suspensions containing RNA beads and two different kinds of cells (APCs and Jurkat reporter cells) were added to each of the two water-phase insets of the droplet-generating chip. RNA-beads with the density of 6,000/μL were mixed into two cell phases (cell density: 8,000/μL), which resulted in about 95.5% of the droplets containing the cell and RNA-bead. As shown on the right of Figure 2A, we calculated the diameter of 1408 generated droplets and found that their average droplet diameter was 89.48 ± 4.43 μm, while the coefficient of variation (CV) was 4.95%. A smaller CV reflected better uniformity of the droplet. Thus, the generated droplets had good uniformity.

**Figure 2.**
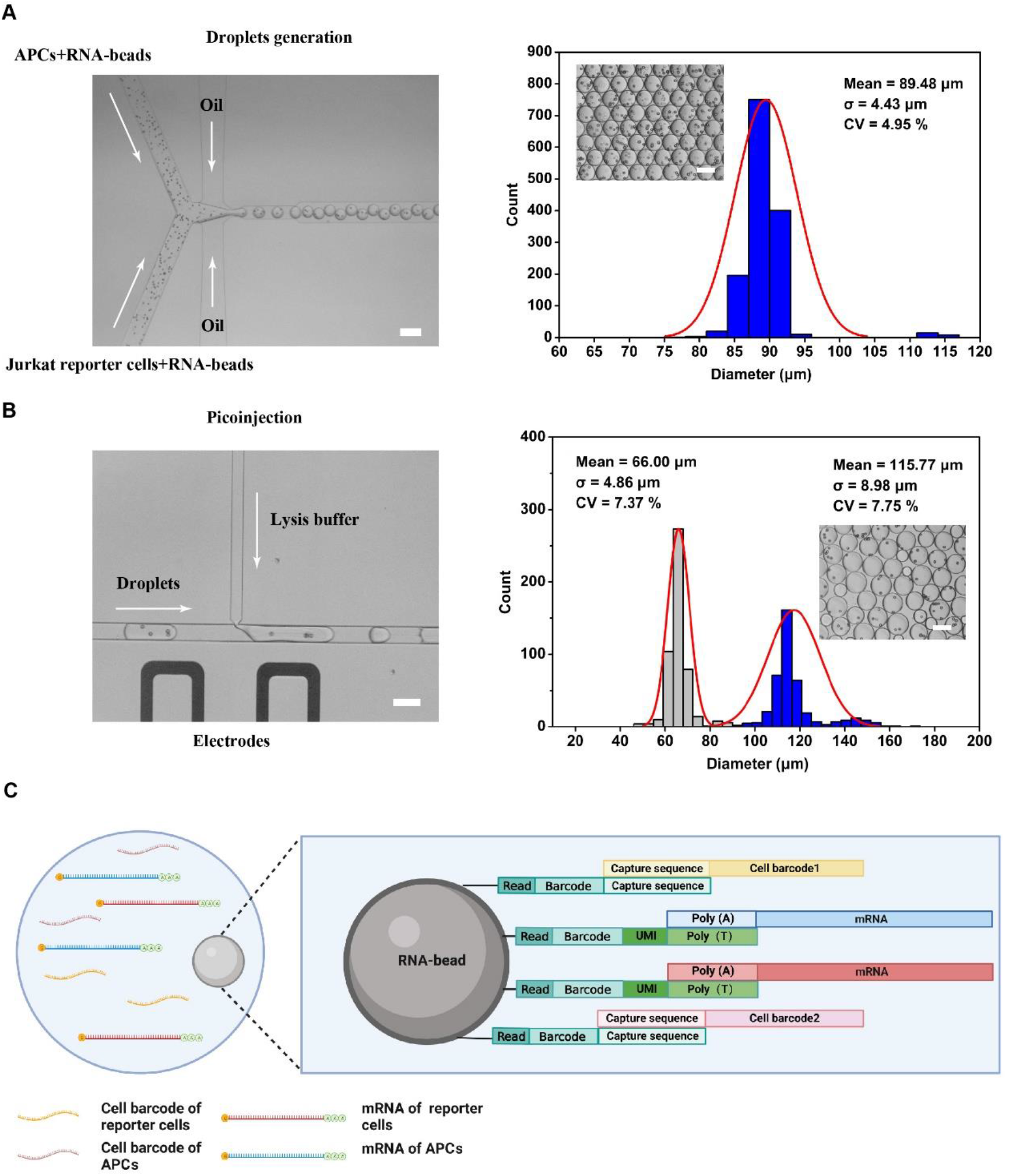
Droplet manipulation and nucleic acid capture. (A) Image of the droplet generation (left). Scale bar represents 200 μm. The right figure shows the diameter distribution of the generated droplets (inset, scale bar represents 100 μm). N=1408. (B) The lysis buffer was picoinjected into the droplet in the manner shown on the left. Scale bar represents 100 μm. The histogram shows the diameter distribution of 890 droplets (inset, scale bar represents 100 μm). (C) The diagram shows nucleic acid (cell barcodes and mRNA) captured on an RNA bead after cell lysis.

After the droplets had been generated, they were collected into a collection tube, which was then placed in an incubator at 37 °C for 4 h. Similar to previous work (Droplet Microfluidics Enables Tracing of Target Cells at the Single-Cell Transcriptome Resolution), we used a pipette to transfer the droplets into a tip, and then inserted the tip into the droplet inlet of the picoinjection chip, which was driven by quickly pulling the syringe from 17.5 to 20 mL. Then, the lysis buffer was injected into the droplet at an applied voltage of 200 V (Figure 2B, left), achieving the capture of cell barcodes and mRNA by RNA beads while retaining the paired relationship of cells in the droplet. The data on droplet diameter as shown in Figure 2B (right) indicated that these droplets were divided into two groups: one comprising small droplets of lysis buffer with a diameter of 66.00 ± 4.86 μm and the other large droplets with a diameter of 115.77 ± 8.98 μm after injection. The injected droplets were then incubated at room temperature for 40 min to achieve cell lysis. A schematic diagram of the nucleic acid (cell barcodes and mRNA) captured on the RNA beads is shown in Figure 2C. Two kinds of sequences were attached to the surface of RNA beads: One was a DNA sequence with a poly-T end for capturing mRNA and the other was a DNA fragment containing the capture sequence for specifically capturing cell barcodes. Finally, the droplets were demulsified and the RNA beads were recovered in accordance with the experimental procedure of single-cell sequencing of the MGI.

### Jurkat reporter system can be used for functional TCR screening

The Jurkat cell line has been widely used to study T-cell activation in response to various antigens in vitro^47–52^. Jurkat reporter cells were previously optimized^49,51^ to have a low signal background and high sensitivity for functional evaluation. Endogenous *b2m, TCRa*, and *TCRb* of Jurkat reporter cells were knocked out and CD8, CD2-CD28, and NF-κB-mCherry were overexpressed in them in our previous experiment (data not shown). The MART-1 melanoma antigen and NY-ESO-1 antigen (Table S2) presented by the class I MHC protein HLA-A*0201 were recognized by TCR clone types DMF5 and 1G4 (Table S1), respectively. They are well-characterized TCR and antigen that have been used in clinical trials of gene-modified T cells. The TCR-DMF5 and TCR-1G4 were overexpressed in Jurkat reporter cells by Sleeping Beauty transposition through electroporation, and then Jurkat reporter TCR cell clones of which at least 90% were expressing TCR were obtained (Figure 3A).

**Fig. 3.**
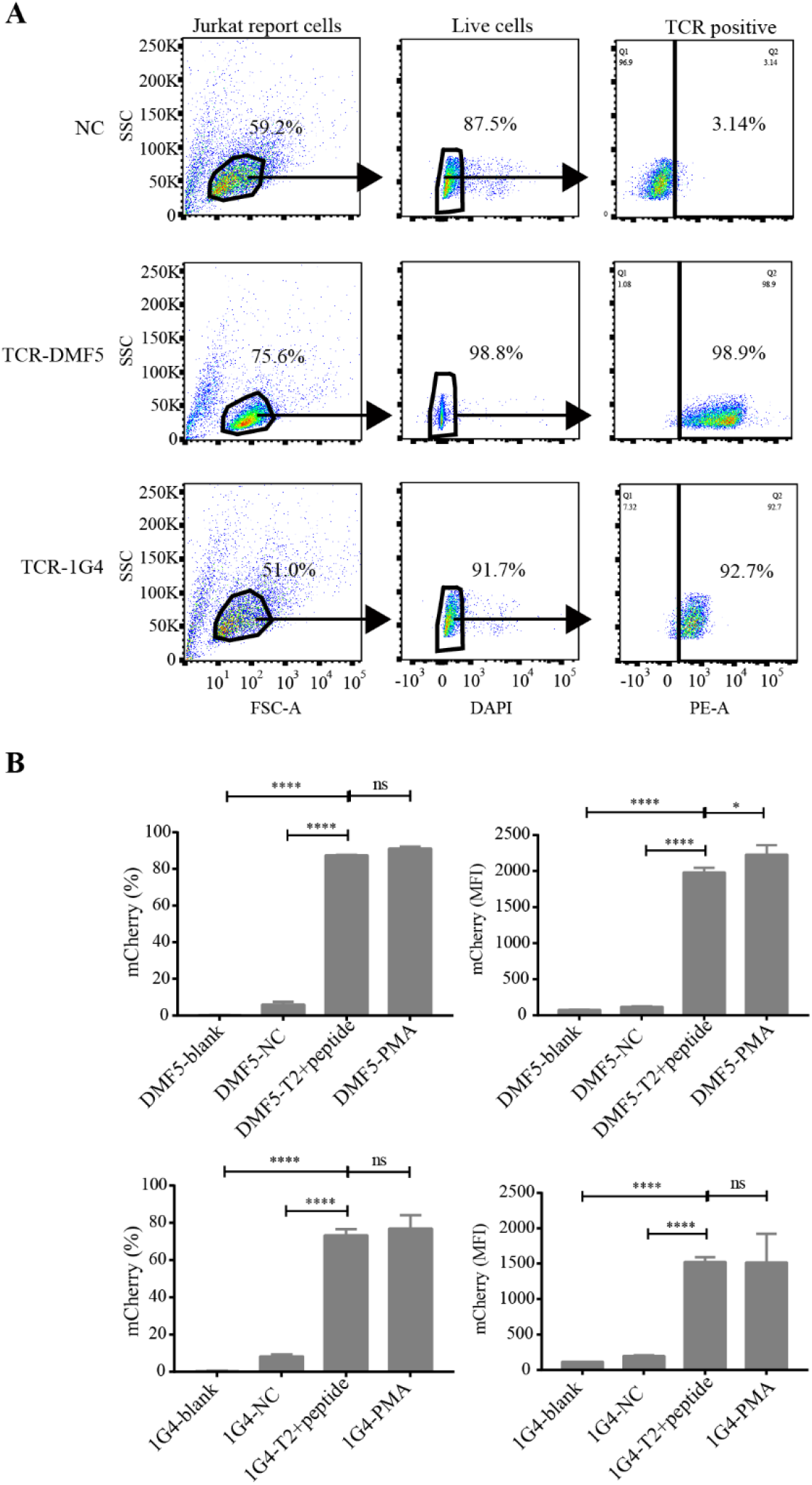
Functional verification of the Jurkat reporter system. (A) The expression of TCR-1G4/TCR-DMF5 in Jurkat reporter cells analyzed by FACS. (B) Validation of Jurkat reporter system stimulated by peptide-pulsed APCs or PMA analyzed by FACS. DMF5-blank group is Jurkat reporter TCR-DMF5 alone without activation, DMF5-NC group is Jurkat reporter TCR-DMF5 cocultured with NY-ESO-1-pulsed T2-HLA-A*0201 cells, DMF5-T2+peptide group is Jurkat reporter TCR-DMF5 cocultured with MART-1-pulsed T2-HLA-A*0201 cells, and DMF5-PMA group is Jurkat reporter TCR-DMF5 alone with PMA stimulation. The 1G4-blank group is Jurkat reporter TCR-1G4 alone without activation, 1G4-NC group is Jurkat reporter TCR-1G4 cocultured with MART-1-pulsed T2-HLA-A*0201 cells, 1G4-T2+peptide group is Jurkat reporter TCR-1G4 cocultured with NY-ESO-1-pulsed T2-HLA-A*0201 cells, and 1G4-PMA group is Jurkat reporter TCR-1G4 alone with PMA stimulation. *p < 0.05, Student’s t test. Data represent mean ± SD, n = 3.

TCR binds its cognate pMHC to activate the transcription factors NFAT and NF-κB, leading to cytokine production in primary T cells^56^. To evaluate the Jurkat reporter system for functional TCR screening, we used T2-HLA-A*0201 cells pulsed with 10 μg/ml MART-1 or NY-ESO-1 peptide coculture with Jurkat reporter TCR-DMF5 and Jurkat reporter TCR-1G4. Upon PMA stimulation or specific peptide pulsing for APC stimulation, over 80% of mCherry fluorescence was detected for both the Jurkat reporter TCR-DMF5 and the Jurkat reporter TCR-1G4, and the mean fluorescence intensity (MFI) of the mCherry fluorescence was at least 1500 (Figures 3B, S1, and S2). For the Jurkat reporter TCR without stimulation or stimulation by negative peptide pulsing of APC, the rate of mCherry fluorescence or MFI was significantly lower than for the specific stimulation (Figures 3B, S1, and S2). Therefore, the Jurkat reporter cells could be used for functional TCR screening with a low signal background and high sensitivity.

### The Jurkat reporter system combined with droplet microfluidics can be used for high-throughput screening of functional antigens and their specific TCRs

To demonstrate that the high-throughput screening platform can determine the cell pairing in each microfluidic droplet by analyzing the DNA barcodes, we constructed Jurkat cells, 293T cells, and K562 cells labeled with unique DNA barcodes to perform a cell-mixing experiment (Figures 4 and S4). Two samples each of Jurkat cells and K562 cells (or HEK293T cells) carried unique DNA barcodes, which were then added to the two water-phase inlets of the droplet-generating chip. In the Jurkat and K562 cell-mixing experiment, 940 cell barcodes were obtained and at least 2000 transcripts were detected. Moreover, in the Jurkat and 293T cell-mixing experiment, 2228 cell DNA barcodes were obtained and at least 2800 transcripts were detected. These cells were successfully assigned to their cell pairing composition using the DNA barcode. Those genes whose expression was enhanced in K562 cells (*BLVRB* and *FAM83A)* and in Jurkat T cells (*CD2* and *CD3D*) exhibited striking expression distribution within K562 cells and Jurkat cells, respectively, which were assigned by DNA barcode and analyzed through volcano plot and UMAP plot (Wilcoxon test, Figure 4A, B). In addition, the genes whose expression was enhanced in 293T cells were highly expressed in 293T cells assigned by the DNA barcode to type 1, as analyzed by volcano plot and UMAP plot (Wilcoxon test, Figure S4A, B). DEGs between the Jurkat cells, K562 cells, and 293T cells were also identified, and those genes acting as exclusive markers of cell types were revealed in a heatmap and dot plot (Figure 4C, D, and Figure S4C, D). These results suggested that the high-throughput screening platform can determine the cell pairing in each microfluidic droplet by analyzing the DNA barcodes with high accuracy and specificity without cell-type-specific bias and with no effect on the gene expression profiles.

**Fig. 4.**
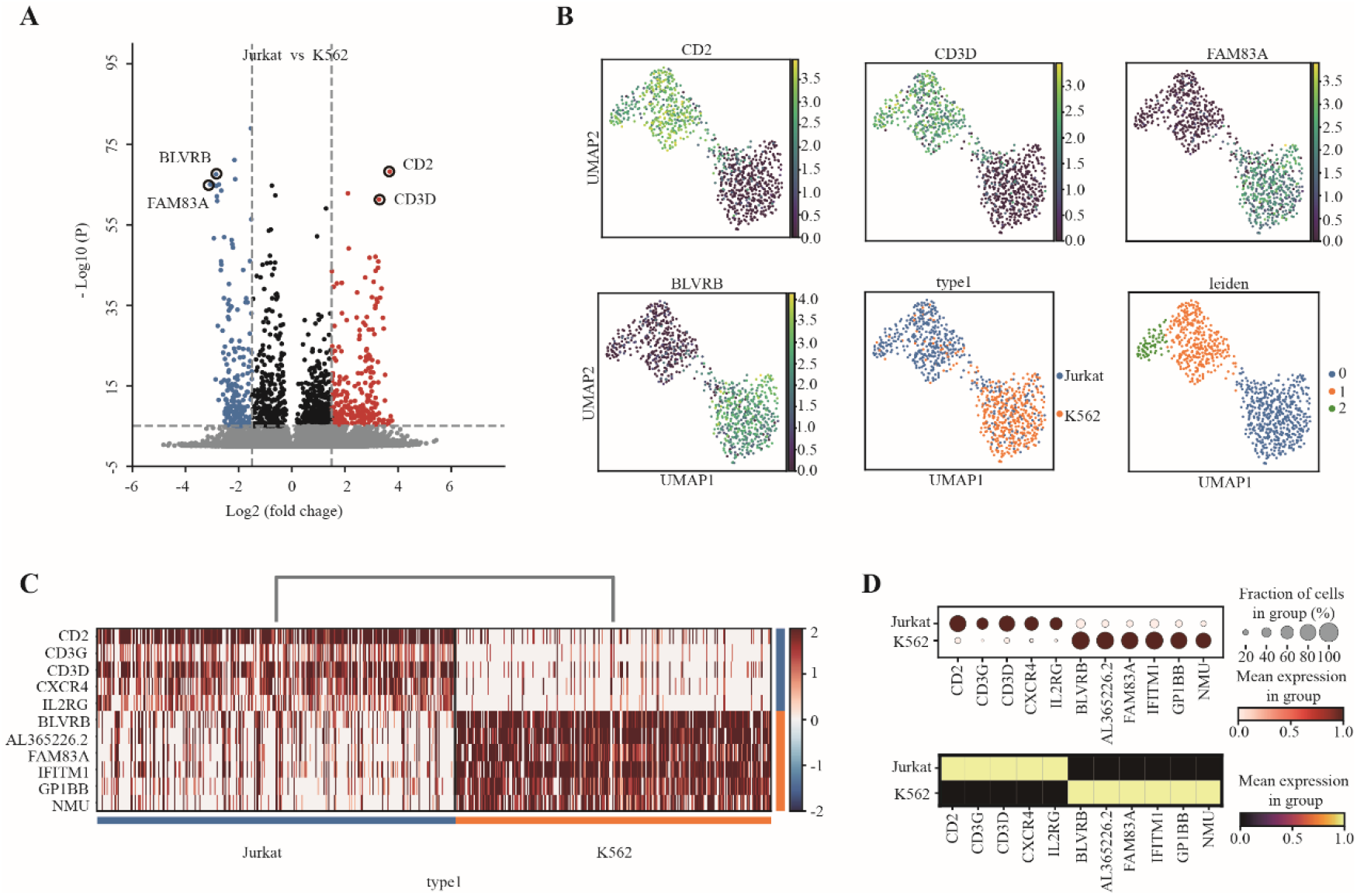
Validation of the screening platform based on the Jurkat reporter combined with droplet microfluidics. A) Volcano plot displaying classification markers of Jurkat cells compared with K562 cells. Genes that have a P value smaller than 0.05 and an absolute value of log(fold change) larger than 1.5 are considered significant. Upregulated genes are colored red, downregulated genes are colored blue, and insignificant genes are colored gray and black. B) UMAP of classification markers in Jurkat reporter cells and K562 cells. C) Heatmap of classification markers in Jurkat reporter cells and K562 cells. D) Matrix plot and dot plot show the classification markers in Jurkat reporter cells and K562 cells. The color density indicates the average expression of a given gene; Wilcoxon test.

Next, we assessed whether the high-throughput screening platform could screen the PMA-stimulated and untreated Jurkat reporter cells. The PMA-stimulated and untreated Jurkat reporter cells labeled with a unique DNA barcode were then added to the phase inlets of the droplet-generating chip and subjected to scRNA-seq. These PMA-stimulated and untreated Jurkat reporter cells were also assigned to their cell origins using the DNA barcode, with the Jurkat T-cell-activation-related genes *CD69, CD82, CD83, DUSP2*, and *ANXA1* being upregulated in PMA-stimulated Jurkat reporter cells, and consistent with the transcription expression of *mCherry* (Figure 5A). There was significantly high expression of Jurkat T-cell-activation-related genes in the PMA-stimulated than in untreated Jurkat reporter cells as analyzed by the heatmap and dot plot (Figure 5B, C). These results suggested that the high-throughput screening platform can determine the cell pairing in each microfluidic droplet by analyzing the DNA barcodes, and screen PMA-stimulated and untreated Jurkat reporter cells through their gene expression.

**Fig. 5.**
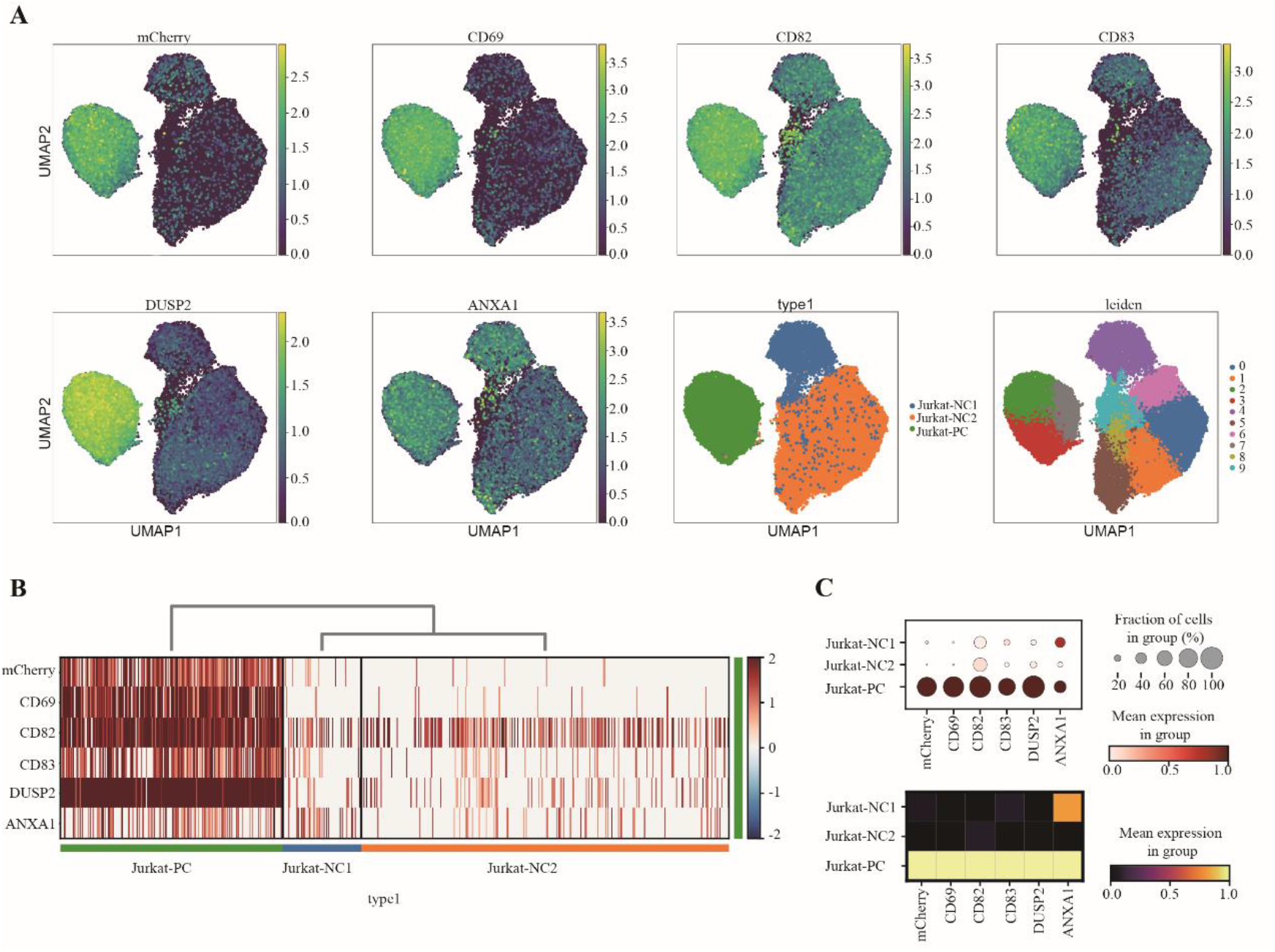
Validation of the screening platform by screening the PMA-stimulated Jurkat reporter cells combined with droplet microfluidics. A) UMAP of T-cell-activation-related DEGs in Jurkat reporter cells before/after PMA stimulation. Cell types Jurkat-NC1 and Jurkat-NC2 were before stimulation. Cell type Jurkat-PC was after PMA stimulation. B) Heatmap of T-cell-activation-related DEGs in each cell type. C) Matrix plot and dot plot show the relative expression of the T-cell-activation-related DEGs in each cell type. Jurkat-PC cells, n = 14,867; Jurkat-NC1, n = 9995; Jurkat-NC2, n = 19,634; Wilcoxon test.

We also validated that the high-throughput screening platform could be used for the high-throughput screening of tumor antigens and their specific functional TCRs. The Jurkat reporter TCR-DMF5 and Jurkat reporter TCR-1G4 were labeled with a unique DNA barcode, as were the APCs (K562-HLA-A*0201) pulsed with MART-1 or NY-ESO-1 peptide. These two Jurkat reporter TCR cells were pooled and then added to the one water-phase inlet of the droplet-generating chip, while these two peptide-pulsed APCs were pooled and added to the other water-phase inlet. Following an experiment involving co-culture, picoinjection, and scRNA-seq, the origins of these cell pairings in droplets were assigned to cell type positive 1 (cell pairing: Jurkat reporter TCR-1G4 with APC-NY-ESO), positive 2 (cell pairing: Jurkat reporter TCR-DMF5 with APC-MART-1), and negative, according to the unique DNA barcodes (Figure 6A). The Jurkat T-cell-activation-related genes *CD69, CD82, CD83, DUSP2, ANXA1*, and *mCherry* were upregulated in cell pairing positive 1 and positive 2 but were downregulated in cell pairing negative, as analyzed by the UMAP plot and dot plot (Figure 6A, B). These results revealed that the high-throughput screening platform can screen the TCR–pMHC-specific stimulated Jurkat reporter cells through the DNA barcode and gene expression. This proof-of-principle study demonstrates that the high-throughput platform can be used for simultaneous, unbiased, library-on-library screening of antigens and their functional TCRs. This platform can also be used to screen the cross-reactivity and off-target recognition of candidate antigens and TCRs in libraries before clinical application to promote tumor immunotherapy.

**Fig. 6.**
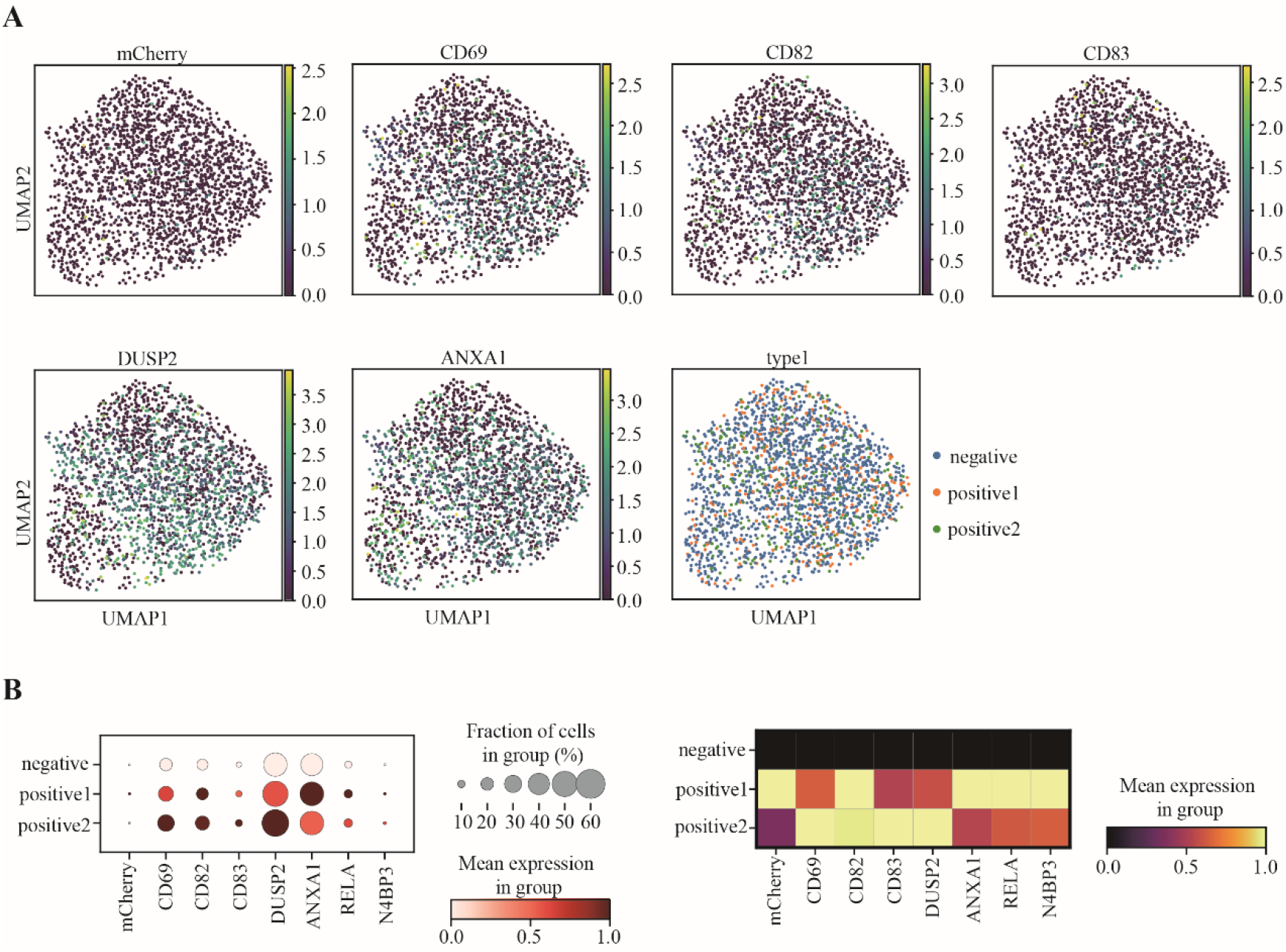
Jurkat reporter system combined with droplet microfluidics can be used for high-throughput screening of tumor antigens and their specific functional TCRs. A) UMAP of T-cell-activation-related DEGs in Jurkat reporter cells before/after PMA. Cell type negative is Jurkat reporter TCR alone, APC alone, Jurkat reporter TCR-DMF5 with APC-NY-ESO, or Jurkat reporter TCR-1G4 with APC-MART-1. Cell type positive 1 is Jurkat reporter TCR-1G4 with APC-NY-ESO. Cell type positive 2 is Jurkat reporter TCR-DMF5 with APC-MART-1. Cell pairings containing all four kinds cells were abandoned. B) Matrix plot and dot plot show the relative expression of the T-cell-activation-related DEGs in each cell type. F) Violin plot shows the T-cell-activation-related DEGs in each cell type. Negative, n = 2142; positive 1, n =449; positive 2, n = 407; Wilcoxon test.

## Discussion

We have developed a high-throughput screening platform, which is combined with a Jurkat reporter system, DNA barcoding, and droplet microfluidics to enable the library-on-library screening of functional antigens and their specific TCR. Jurkat reporter TCR cells could be used for functional TCR screening with a low signal background and high sensitivity (Figure 3). By replacing primary T cells, they could also significantly decrease the cost of functional TCR screening. By using droplet microfluidics and picoinjection, unique DNA barcode-labeled Jurkat reporter TCR cell mix and APC mix were encapsulated in the same droplet. Then, lysis buffer was injected into the droplet to release the DNA barcode and mRNA without breaking the cell pairings in the droplet. Upon analyzing the DNA barcodes and T-cell-activation-pathway-related genes combined with *mCherry* expression, the cell pairing in each droplet and the specific recognition relationship between the antigen and its functional TCR were able to be interpreted (Figures 5 and 6). The high-throughput screening platform provided a powerful tool for the identification of tumor antigens and specific TCRs simultaneously, which could significantly decrease the time required for screening functional antigens and specific TCRs or screening cross-reactivity and off-target recognition.

To date, some high-throughput methods for T-cell antigen identification have been established. For TCR and pMHC affinity-based assays such as pMHC multimers^26,27^, yeast display^28,29^, TCR-pMHC interaction Koff-Rate^30^, YAMTAD^31^, and dextramer-TCR binding assays^32^, these could identify high-affinity TCRs but would significantly increase the risk of cross-reactivity and off-target recognition^34,35^. Moreover, affinity might not be a reliable indicator of T-cell response^37,38^, and TCR-pMHC interaction may be related to high binding affinity but not specific stimulatory interaction. For T-cell function- or TCR active signaling pathway-based assays, such as trygocytosis^39,40^, T-scan^41^, “catch bound” ^42^, cytokine-capturing^43^, or combining single-cell droplet microfluidics with TCR active signaling pathway^44,45^, challenges regarding limited throughput still remain. They cannot achieve unbiased screening of functional TCR and antigen simultaneously when there are candidate TCR libraries and peptide libraries, and thus evaluating the interaction between these two libraries would be very time-consuming^21^.

Our high-throughput screening platform using the active signaling pathway in Jurkat reporter TCR cells to screen the functional TCR could be more credible and decrease the cost of screening. Combined with droplet microfluidics and picoinjection, the unique DNA barcode-labeled Jurkat reporter TCR cell mix and APC mix were encapsulated in a droplet and the DNA barcode and mRNA were then released without breaking the cell pairing in the droplet. With the help of scRNA-seq, unique DNA barcodes and T-cell-activation-pathway-related genes combined with *mCherry* expression could be used to interpret the cell pairing in each droplet and the specific recognition relationship between these candidate antigens and their functional TCRs in unbiased library-on-library screening. Moreover, the high-throughput platform could also be used to screen the cross-reactivity and off-target recognition of candidate antigens and TCRs before clinical application to promote tumor immunotherapy. In further research, the high-throughput screening platform would be modified by combining the Jurkat reporter APCs, DNA barcoding, and droplet microfluidics with full-length transcriptome sequencing methods to achieve the major breakthrough of high-throughput screening of neoantigen-specific TCRs in peripheral blood of clinical cancer patients.

## Limitations of the study

Our study is limited by the lack of additional functional tumor antigens and their specific TCRs because these two functional antigens and TCRs may be insufficient to validate the high-throughput nature of the screening platform. At present, our team is accumulating functional neoantigen and TCR data, which will be used for the verification of high-throughput screening platforms. Our studies are also limited by the low efficiency of DNA barcode labeling (∼50%), which may result in false negative findings for some cell pairings, while having no effect on the cell pairings assigned as positive. Our team continues to work to optimize DNA labeling efficiency. Gene Ontology (GO) analysis also showed no enrichment of the T-cell activation pathway when analyzing the tumor antigens and their specific functional TCRs (data not shown), which may be due to the difference in activation of Jurkat T cells and primary T cells.

## Acknowledgments

We acknowledge financial support from the National Natural Science Foundation of China (Grant No. 82202040), National Key Research and Development Program of China (Grant No. 2021YFF1200500), and Guangdong Basic and Applied Basic Research Foundation (Grant No. 2021A1515110459). This work was supported by China National GeneBank (CNGB), Guangdong Provincial Key Laboratory of Human Disease Genomics (2020B1212070028), Shenzhen Key Laboratory of Genomics (CXB200903110066A), Shenzhen Key Laboratory of Single-Cell Omics (ZDSYS20190902093613831), Guangdong Provincial Key Laboratory of Genome Read and Write (2017B030301011).

## Supplementary data

Supplementary data to this article can be found online.

## STAR METHODS

## KEY RESOURCES TABLE

| REAGENT or RESOURCE | SOURCE | IDENTIFIER |
| --- | --- | --- |
| Antibodies |  |  |
| PE Hamster Anti-Mouse TCR $\beta$ Chain | BD Biosciences | Cat#553172 |
| DAPI Solution | BD | Cat#564907 |
| Chemicals, peptides, and recombinant proteins |  |  |
| MART-1 peptide | GenScript | N/A |
| NY-ESO-1 peptide | GenScript | N/A |
| pCMV(CAT)T7-SB100X plasmid | Addgene | Cat#34879 |
| pT2/HB plasmid | Addgene | Cat#26557 |
| TCR-DMF5 DNA fragment | Beijing Liuhe BGI Co., Ltd. | N/A |
| TCR-1G4 DNA fragment | Beijing Liuhe BGI Co., Ltd. | N/A |
| pT2/HB-HLA-A*1101 DNA fragment | Beijing Liuhe BGI Co., Ltd. | N/A |
| pT2/HB-HLA-A*0201 DNA fragment | Beijing Liuhe BGI Co., Ltd. | N/A |
| Critical commercial assays |  |  |
| Electroporation kits | Celetrix | Cat#1201 |
| DNBelab C4 kit | MGI | Cat#940-00050 |
| Deposited data |  |  |
| fetal bovine serum | HYCLONE | Cat#SH30084.03HI |
| RPMI 1640 | HYCLONE | Cat# SH30809.01 |
| IMDM | GIBCO | Cat#12440053 |
| DMEM | INVITROGEN | Cat#C11995500BT |
| Penicillin-Streptomycin (10,000 U/mL) | GIBCO | Cat#15140122 |
| dimethyl sulfoxide (DMSO) | SIGMA | Cat#D5879-500ML |
| OPTI-MEM | INVITROGEN | Cat#11058-021 |
| PMA+Ionomycin (PMA) | Dakewe Biotech Co., Ltd. | Cat#2030421 |
| Puromycin | INVIVOGEN | Cat#ant-pr-1 |
| PBS | LIFE TECHNOLOGIES | Cat#14040-133 |
| Lipofectamine 3000 | THERMO | Cat#L3000015 |
| oxygen plasma | Harrick Plasma | Cat#PDC-002 |
| PDMS | DOW CORNING | Cat#DC059 |
| SU-8 molds | HICOMP MicroTech(Suzhou)<br>Company | N/A |
| Ficoll PM-400 | SIGMA | Cat#F4375-25G |
| oil | Bio-Rad | Cat#1864005 |
| 10% Pluronic F68 | SIGMA | Cat#P5556 |
| Experimental models: Cell lines |  |  |
| T2 cell | ATCC | RRID: CVCL_2211 |
| K562 cell | ATCC | RRID: CVCL_0004 |
| HEK293T | ATCC | RRID: CVCL_0063 |
| Jurkat E6-1 cell | ATCC | RRID: CVCL_0367 |
| Jurkat reporter cell | This paper | N/A |
| Jurkat reporter TCR cell | This paper | N/A |
| Oligonucleotides |  |  |
| DNA barcode (ssDNA) | Beijing Liuhe BGI Co., Ltd. | N/A |
| Software and algorithms |  |  |
| FlowJo (10.5.3) | Tree Star/BD Biosciences | https://docs.flowjo.com/flowjo/getting-acquainted/10-5-release-notes/10-5-3-release-notes/10-5-3-exhaustive-release-notes/ |
| Prism (GraphPad) | Graphpad by Dotmatics | https://www.graphpad.com/scientific-software/prism/ |
| Seurat | New York Genome Center, Satija Lab | https://github.com/satijalab/seurat |
| Data files for single-cell RNA sequencing (raw data) | This paper | **CNP0003756** |

## EXPERIMENTAL MODEL AND SUBJECT DETAILS

### Materials and methods

#### Peptide synthesis

Tumor-associated/specific peptides (Table S1) were synthesized by GenScript (Nanjing, China), the purity of which exceeded 99.0%. Peptides were dissolved with 100% dimethyl sulfoxide (DMSO; Sigma-Aldrich, USA) at a concentration of 10 mg/mL and stored at −20°C.

#### Cell culture

T2 cells (RRID: CVCL_2211), K562 cells (RRID: CVCL_0004), and HEK293T cells (RRID: CVCL_0063) were purchased from American Type Culture Collection. In the Jurkat reporter cell line based on the Jurkat E6-1 cell line (RRID: CVCL_0367), endogenous β2m, and TCRα and β chains were knocked out, followed by the overexpression of CD8, CD2-CD28, and NF-κB-mCherry, as previously described^51^. Jurkat cells and K562 cells were cultured in RPMI 1640 (Gibco, USA) supplemented with 10% (v/v) fetal bovine serum (FBS; HYCLONE, USA) containing 1% penicillin–streptomycin (P/S) at 37 °C with 5% CO_2_. T2 cells were cultured in IMDM (Gibco, USA) supplemented with 10% (v/v) FBS and 1% P/S at 37 °C with 5% CO_2_. HEK293T cells were cultured in DMEM (Gibco, USA) supplemented with 10% (v/v) FBS and 1% P/S at 37 °C with 5% CO_2_.

#### Generation of candidate TCR-overexpressing Jurkat reporter cells

Overexpression of the candidate TCRs was achieved using Sleeping Beauty transposition with the pCMV(CAT)T7-SB100X plasmid obtained from Addgene (Plasmid #34879). These candidate TCRs (Table S2) were synthesized by Beijing Liuhe BGI Co., Ltd. (Beijing, China), and then cloned in the pT2/HB plasmid obtained from Addgene (Plasmid #26557), into which the hPGK promoter (human phosphoglycerate kinase 1 promoter) had been inserted. The variable regions of these candidate TCRs were fused with murine TCRα and β2 constant regions. These TCRα and β chains were linked by a P2A self-cleaving peptide. In the electroporation process, approximately 6×10^6^ Jurkat reporter cells were washed three times with OPTI-MEM (Gibco, USA) and resuspended with 120 μl of electroporation buffer (Celetrix, Germany). Subsequently, 12 µg of pCMV(CAT)T7-SB100X plasmid and 24 µg of pT2/HB-TCR plasmid were electroporated into the cells using the Electroporator SP100 (Celetrix, Germany) at 795 V for 20 ms. Following this electroporation, these cells were immediately placed in pre-warmed culture medium at 37°C and 5% CO _2_. Finally, the Jurkat reporter TCR cells were obtained by screening for puromycin resistance 5 days after the electroporation.

#### Generation of HLA-A*1101/HLA-A*0201-overexpressing K562 cells

The HLA-A*1101 gene fragment (GenBank: D16841.1) and HLA-A*0201 gene fragment (GenBank: U02935.2) were synthesized at Beijing Liuhe BGI Co., Ltd. (Beijing, China), and then cloned in the modified pT2/HB plasmid. In the electroporation process, approximately 6×10^6^ K562 cells were washed three times with OPTI-MEM (Gibco, USA) and resuspended with 120 μl of electroporation buffer (Celetrix, Germany). Next, 12 µg of pCMV(CAT)T7-SB100X plasmid and 24 µg of pT2/HB-HLA-A*1101/ pT2/HB-HLA-A*0201 plasmid were electroporated into the cells using the Electroporator SP100 (Celetrix, Germany) at 795 V for 20 ms. Following this electroporation, these cells were immediately placed in pre-warmed culture medium at 37°C and 5% CO _2_. Finally, the K562-HLA-A*1101 and K562-HLA-A*0201 cells were obtained by screening for puromycin resistance 5 days after the electroporation.

#### Flow cytometry

To analyze the Jurkat reporter TCR cells, Jurkat reporter TCR-1G4 and Jurkat reporter TCR-DMF5 were labeled with PE Hamster Anti-Mouse TCR β Chain (BD Biosciences, USA). Staining of cell surface proteins was performed in flow buffer [phosphate-buffered saline (PBS) + 0.5% FBS] at 4°C in the dark for 30 min. After staining, cells were washed and resuspended in flow buffer. FACS analysis was performed using a FACS Arial II flow cytometer (Becton Dickinson, USA). Flow cytometry data were analyzed using software FlowJo (FlowJo, USA).

#### Mean fluorescence intensity analysis

Suspensions of Jurkat reporter TCR-1G4 and Jurkat reporter TCR-DMF5 cells (5×10^4^/100 μl/well) were added to each well of 96-well plates. The APCs T2-HLA-A*0201 were pulsed with MART-1 and NY-ESO-1 at a concentration of 10 μg/ml for 24 h, after which they were added (at 5×10^4^/100 μl/well) to each well at a ratio of 1:1. In all groups, the experiments were performed in triplicate. PMA (cat. no. 2030421; Dakewe, China) stimulation was applied to the cells at a concentration of 5 *μ*g/ml. After 48 h of incubation at 37°C and 5% CO _2_, the cells were washed and resuspended in flow buffer. FACS analysis was performed using a FACS Arial II flow cytometer (Becton Dickinson, USA). Flow cytometry data were analyzed using FlowJo (FlowJo, USA).

#### ssDNA design and transfection

The DNA barcode (ssDNA) contained a unique sample barcode (52 nucleotides), an amplification handle (34 nucleotides), and a capture tail (14 nucleotides). The full ssDNA sequence is as follows: 5′-TTTTGTCTTCCTAAGACCGCTTGGCCTCCGA CTT(N)_52_GACGCTGCCGACGA-3′. The amplification handle sequence is “TTTTGTCTTCCTAAGACCGCTTGGCCTCCGACTT,” where the “N” is the unique sample barcode sequence and “GACGCTGCCGACGA” is the capture tail sequence. All ssDNAs (Table S3) were synthesized at Beijing Liuhe BGI Co., Ltd. (Beijing, China), without any modifications. Then, the cells were transfected with ssDNA (80 pmol/ml, 1×10^6^ cells per well) using Lipofectamine 3000 (Life Technologies, USA), in accordance with the manufacturer’s protocol.

#### Microfluidic chip design and fabrication

All microfluidic chips were designed using AutoCAD software, including a droplet-generating chip and a picoinjection chip. The SU-8 molds were obtained by standard lithography. The PDMS chips were obtained by pouring mixed polydimethylsiloxane (PDMS, 10:1 base-to-curing agent ratio) onto these SU-8 molds and placing them in an oven at 90°C for 2 h. After curing, we peeled the PDMS from the mold, and obtained chips with channels of 90 or 60 μm in height. The PDMS chips were then cut into the desired shape with a cutter knife, and holes were punched at the inlet and outlet of the chip with differently sized punches. For the parameters for fabricating the droplet-generating chip, please refer to a previous paper^54^. As for the picoinjection chip, Figure S3 shows the design parameters; the diameter of the holes of all inlets and outlets was 1 mm. The two kinds of chips were bound to PDMS substrate or glass substrate by oxygen plasma (Harrick Plasma, PDC-002) treatment, and heated in an oven at 110°C for 30 min. For the picoinjection chip, the electrodes were manufactured by injecting melted low-melting-point solder wires into the electrode microchannels on a hot plate at 90 °C, after which the chip was removed from the hot plate in order to solidify the electrodes.

#### Microscope imaging scRNA-seq

Bright-field images were captured by an Olympus microscope (IX71; Olympus, Japan) with a high-speed camera (DP26; Olympus, Japan). Meanwhile, images of fluorescently labeled Jurkat reporter TCR cells were acquired using an inverted fluorescence microscope (IX73; Olympus, Japan). Droplets were dropped on a glass slide and imaged with a 10× objective lens of an inverted microscope. Then, these images were automatically quantified by ImageJ software to acquire quantitative results.

#### Single-cell RNA library preparation and sequencing

First, the cells and RNA beads were suspended in working buffer (RPMI 1640 supplemented with 10% FBS, 1% P/S, 0.1% Pluronic F-68, and 6% Ficoll PM400). Different types of suspensions were added to the two water-phase inlets, and oil (1864005; Bio-Rad, CA, USA) was added to the oil-phase inlet. The outlet was connected through a pipe to the entrance of a special collection tube, which was connected to a 30 mL syringe fixed to a three-dimensional printed base^55^. The droplet generation was driven by pulling the syringe from 15 to 20 mL to create negative pressure. Second, after being placed in an incubator at 37°C for 4 h, the droplets were then sucked out with a 200 μL tip, and the tip was inserted into the droplet inlet of the picoinjection chip. The syringe was pulled to provide negative pressure, and the voltage was adjusted to 200 V for the injection of lysis buffer. Third, after the picoinjection, the droplets were incubated for 40 min at room temperature and then demulsified to recover RNA beads. The cDNA library for scRNA-seq was generated using the DNBelab C4 kit. Fourth, all libraries were conducted in preparation for sequencing by DIPSEQ T1 sequencer (MGI). The FASTQ data generated by the sequencer were processed with the PISA pipeline (https://github.com/single-cell-BGI/scRNA-pipe), and a standard format gene expression matrix was generated with reference to the GRCh38 human genome for downstream analysis.

#### Data analysis

The gene expression matrix was processed as input for Seurat software (https://github.com/satijalab/seurat). Genes expressed in less than three cells and cells with less than 200 genes were filtered out. To minimize the batch effect among samples, all samples’ analyses were treated by the reciprocal PCA pipeline (parameter: dims = 20) in Seurat for merging into an integrated dataset. The first 50 principal components (PCs) across highly variable genes (HVGs) were summarized to construct an SNN network, and distance between cells and clusters was calculated by a Louvain algorithm. Then, dimension reduction and the plotting of cells in two-dimensional space were performed with UMAP.

#### Differentially expressed gene (DEG) analyses

Identification of the DEGs among the clusters by the “FindAllMarkers” function in Seurat revealed the heterogeneity among TCR-T cells. Genes with adjusted *p* < 0.05 and fold change > 2 were defined as DEGs.

#### Statistical analysis

Statistical analyses were conducted by GraphPad and Seurat. The significance of differences between two groups was examined by Wilcoxon test, and the *p* values from multiple tests were adjusted by the false discovery rate (FDR) method.

#### Data availability

The data that support the findings of this study have been deposited into CNGB Sequence Archive (CNSA) of China National Genebank Database (CNGBdb) with accession number CNP0003756.

