## supplementary data for "High-throughput screening of functional neo-antigens and their specific TCRs via the Jurkat reporter system combined with droplet microfluidics"

**Table S1 Functional TCR clone types**

| TCR clone types | CDR3 sequence: TRA | CDR3 sequence: TRB | HLA type |
|---|---|---|---|
| DMF5 | CAVNFGGGKLIF | CASSLSFGTEAFF | HLA-A*0201 |
| 1G4 | CAVRPTSGGSYIPTF | CASSYVGNTGELFF | HLA-A*0201 |

**Table S2 Functional peptides**

| Peptide | Sequence | HLA type |
|---|---|---|
| MART-1 | LAGIGILTV | HLA-A*0201 |
| NY-ESO-1 | SLLMWITQC | HLA-A*0201 |

**Table S3 DNA barcode sequence**

| Name | Sequence | Cell | Experiment |
|---|---|---|---|
| Barcode1 | TTTTGTCTTCCTAAGACCGCTTGGCCT<br>CCGACTTNNNNNNNNNNTCTGCGTGT<br>GTGTAACCCTTCCTAACAGCCAANNN<br>NNNNNNNGACGCTGCCGACGA | 293T | 293T vs Jurkat |
| Barcode2 | TTTTGTCTTCCTAAGACCGCTTGGCCT<br>CCGACTTNNNNNNNNNNTCTGCGGAT<br>AATGATGCCTTCCTAACAGCCAANNN<br>NNNNNNNGACGCTGCCGACGA | Jurkat | |
| Barcode3 | TTTTGTCTTCCTAAGACCGCTTGGCCT<br>CCGACTTNNNNNNNNNNTCTGCGCT<br>GATACTTCCCTTCCTAACAGCCAANN<br>NNNNNNNNGACGCTGCCGACGA | Jurkat | |
| Barcode4 | TTTTGTCTTCCTAAGACCGCTTGGCCT<br>CCGACTTNNNNNNNNNNTCTGCGGG<br>CTCCAAGCCCTTCCTAACAGCCAANN<br>NNNNNNNNGACGCTGCCGACGA | Jurkat | |
| Barcode5 | TTTTGTCTTCCTAAGACCGCTTGGCCT<br>CCGACTTNNNNNNNNNNTCTGCGAAT<br>CTACAAGCCTTCCTAACAGCCAANNN<br>NNNNNNNGACGCTGCCGACGA | Jurkat | Jurakt vs K562 |
| Barcode6 | TTTTGTCTTCCTAAGACCGCTTGGCCT<br>CCGACTTNNNNNNNNNNTCTGCGCG<br>AATATAGGCCTTCCTAACAGCCAANN | K562 | |

| | | | |
|---|---|---|---|
| | NNNNNNNNGACGCTGCCGACGA | | |
| Barcode7 | TTTTGTCTTCCTAAGACCGCTTGGCCTCCGACTTNNNNNNNNNNNTCTGCGTACGCCGATTCCTTCCTAACAGCCAANNNNNNNNNNGACGCTGCCGACGA | K562 | |
| Barcode8 | TTTTGTCTTCCTAAGACCGCTTGGCCTCCGACTTNNNNNNNNNNNTCTGCGAAGATTAACCCCTTCCTAACAGCCAANNNNNNNNNNGACGCTGCCGACGA | K562 | |
| Barcode9 | TTTTGTCTTCCTAAGACCGCTTGGCCTCCGACTTNNNNNNNNNNNTCTGCGCTAAGAGTCCCCTTCCTAACAGCCAANNNNNNNNNNGACGCTGCCGACGA | K562 pulsed with NY-ESO-1 | 2 TCRs vs 2 peptides |
| Barcode10 | TTTTGTCTTCCTAAGACCGCTTGGCCTCCGACTTNNNNNNNNNNNTCTGCGTGACGTCCTTCCTTCCTAACAGCCAANNNNNNNNNNGACGCTGCCGACGA | K562 pulsed with MART-1 | |
| Barcode11 | TTTTGTCTTCCTAAGACCGCTTGGCCTCCGACTTNNNNNNNNNNNTCTGCGTTGAAGAAGCCCTTCCTAACAGCCAANNNNNNNNNNGACGCTGCCGACGA | Jurkat-TCR-1G4 | |
| Barcode6 | TTTTGTCTTCCTAAGACCGCTTGGCCTCCGACTTNNNNNNNNNNNTCTGCGCGAATATAGGCCTTCCTAACAGCCAANNNNNNNNNNGACGCTGCCGACGA | Jurkat-TCR-DMF5 | |

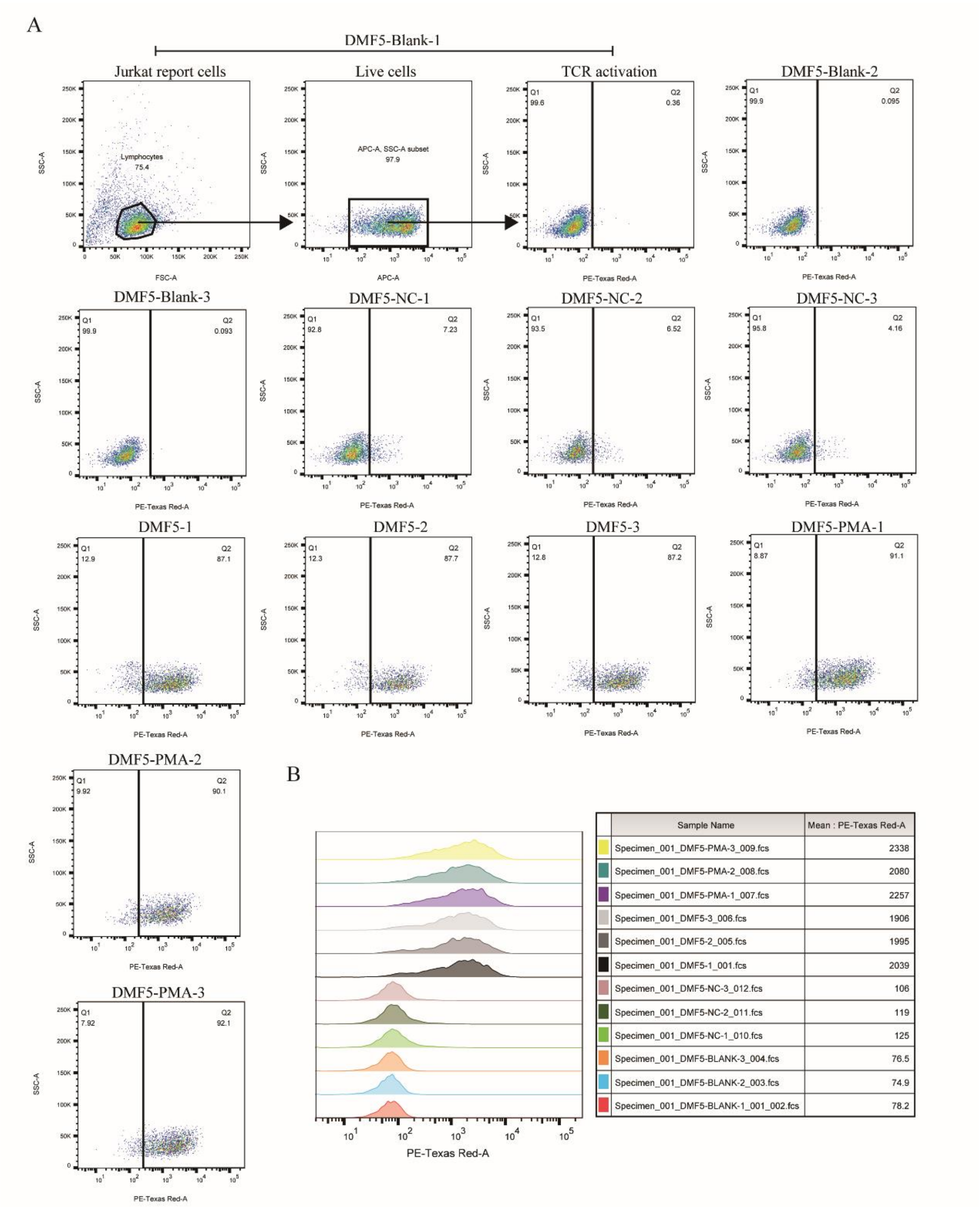


**Fig. S1.** Functional verification of the Jurkat reporter system by TCR-DMF5. (A, B) Validation of the Jurkat reporter system stimulated by peptide-pulsed APCs or PMA analyzed by FACS.

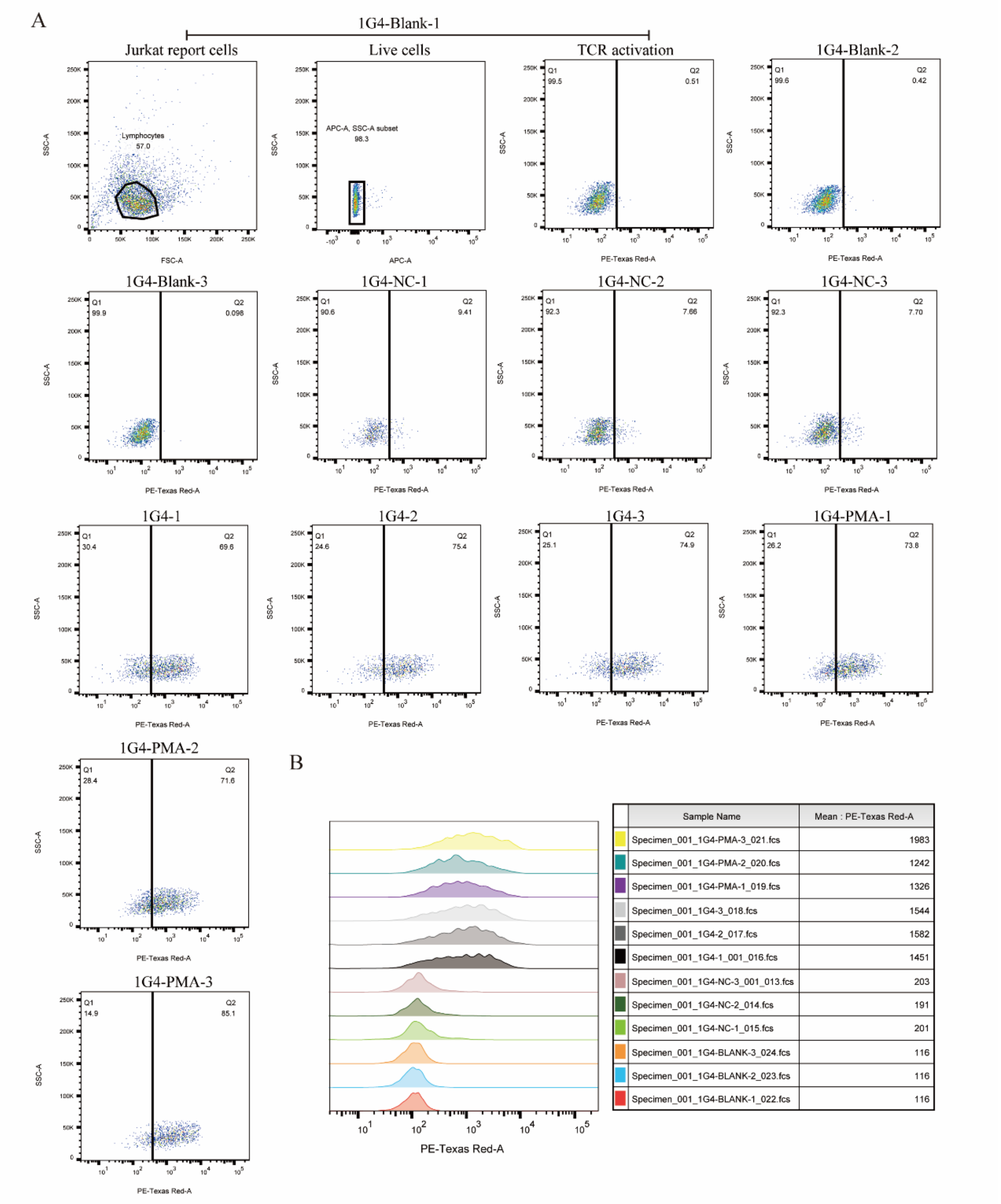


**Fig. S2.** Functional verification of the Jurkat reporter system by TCR-1G4. (A, B) Validation of the Jurkat reporter system stimulated by peptide-pulsed APCs or PMA analyzed by FACS.

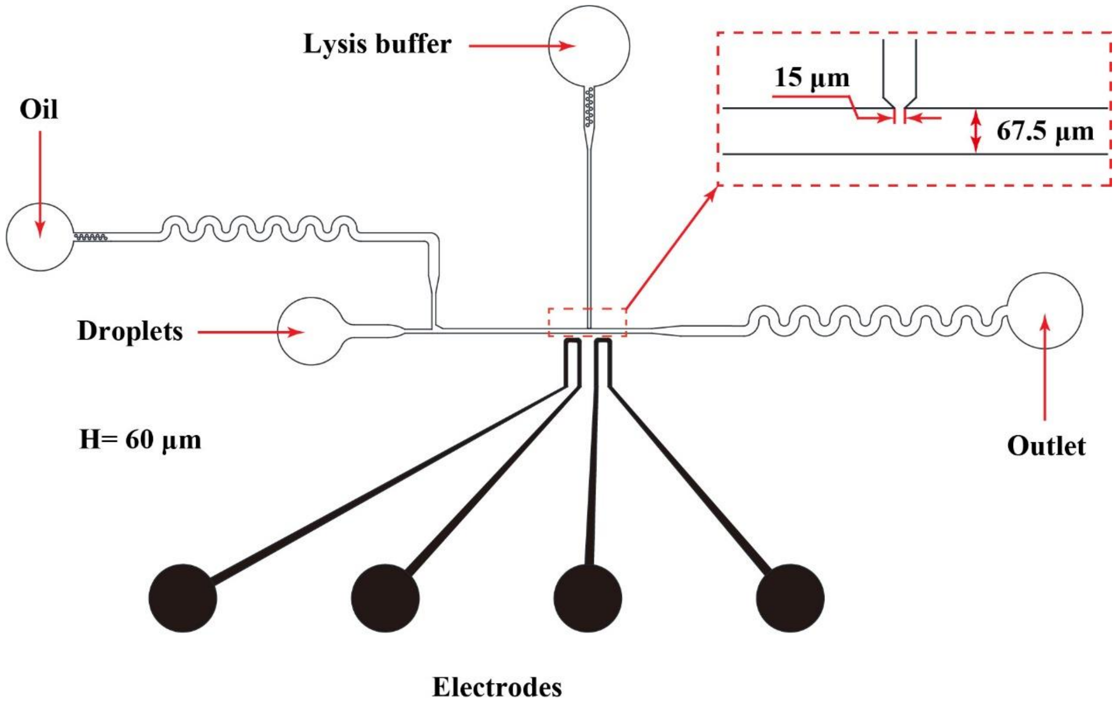


**Fig. S3.** Schematic of droplet picoinjection chip. This chip consisted of a droplet inlet, an oil inlet, an injection phase (lysis buffer) inlet, an outlet, and electrodes. The height of the microfluidic channel was 60 µm.

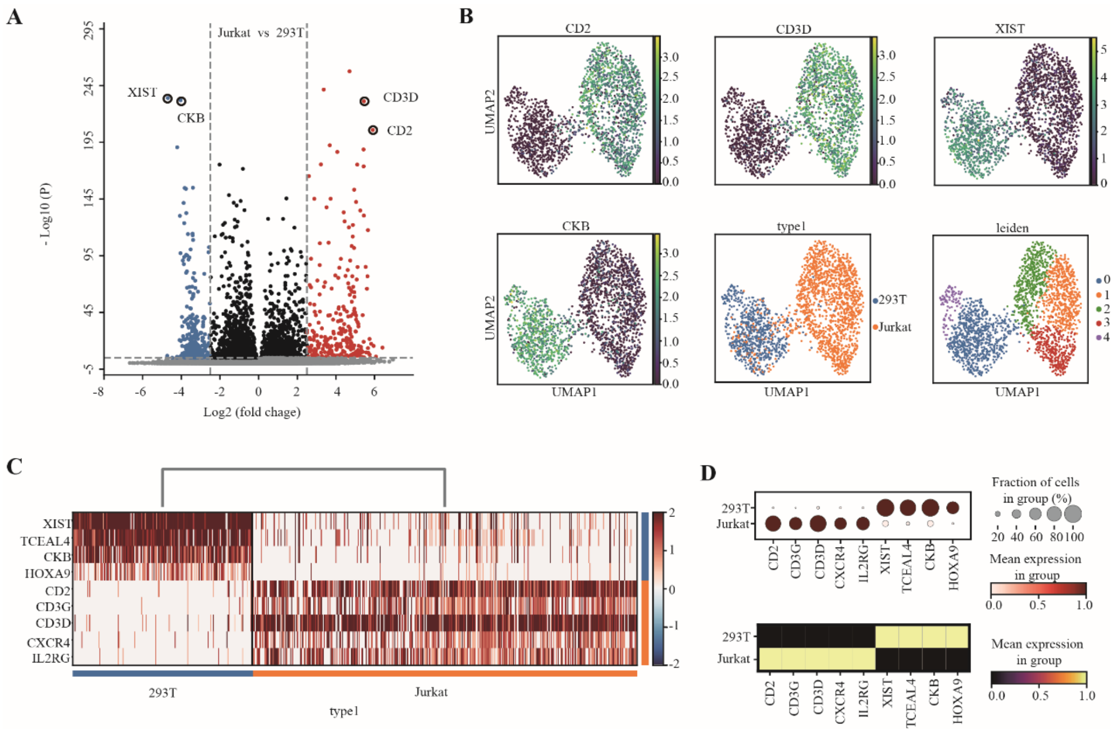


**Fig. S4.** Validation of the screening platform based on the Jurkat reporter combined with droplet microfluidics. A) Volcano plot displaying classification markers of Jurkat cells compared with 293T cells. Genes with a P value smaller than 0.05 and an absolute value of log(fold change) larger than 2.5 are considered significant. Upregulated genes are colored red, downregulated genes are colored blue, and insignificant genes are colored gray and black. B) UMAP of classification markers in Jurkat reporter cells and 293T cells. C) Heatmap of classification markers in Jurkat reporter cells and 293T cells. D) Matrix plot and dot plot show the classification markers in Jurkat reporter cells and 293T cells.
