## Supplementary material for "High-throughput screening of functional neo-antigens and their specific TCRs via the Jurkat reporter system combined with droplet microfluidics": &#20262;&#29702;&#35768;&#21487;

2022041833

### 华大生命伦理审查批件

THE INSTITUTIONAL REVIEW BOARD OF BGI ETHICAL CLEARANCE

批件号 Doc No. BGI-IRB 22069　　有效期至 Expired on 2024/04/14

项目大类 Project Category: 涉及人的生命科学研究Research Involving Humans

项目名 Project Title: 肿瘤免疫治疗信息库建设

负责人 Person in Charge: 刘龙奇　
董旋　

委员会声明 IRB Statement:
・本委员会参照WMA《赫尔辛基宣言》、CIOMS《涉及人的健康相关研究国际伦理指南》（2016）、原卫计委《涉及人的生物医学研究伦理审查办法》（2016）等相关法规、指南标准组成和运作
BGI-IRB is organized and operated complying with WMA Declaration of Helsinki, CIOMS International Ethical Guidelines for Health-related Research Involving Humans 2016 and applicable regulations /guidelines
・本委员会已在广东省深圳市卫健委登记备案
BGI-IRB has registered itself at local authorities

审查方式 Review Conducted: 初始审查Initial Review　　程序 Procedures: 简易（快速）程序Expedited Procedure

意见 Comments:
本项目为华大主导发起的研究项目，结合已有的与中山大学附属第六医院的合作研究项目（原项目已通过华大伦理委员会批准，编号为BGI-IRB 19141“不同分期结直肠癌多组学研究-结直肠癌多维度生命组学图谱绘制”）中的100例肿瘤组织样本和已产生的组学数据进行研究，拟通过质谱技术鉴定新抗原，对肿瘤免疫治疗信息库进行完善。项目所使用的信息均经匿名化处理，数据库访问做好权限限制和安全管理，经委员会审查，意见如下：

决定 Conclusions: 同意Approval　　跟踪审查 Continuing Review: ○ 不适用 N/A　◉ 不晚于 prior to 2023/06/30

注意事项 Reminders:
1.实施前，法规和监管部门有特定要求的，应事先取得相应许可
Specific permits required by applicable regulations should be obtained prior to implementation
2.实施过程及必要文档须与审查通过方案一致，并遵循科研诚信准则
Implementation and achievement should be consistent with the approved proposal and comply with principles of research integrity
3.实施中，如需改变方案并可能显著改变整体风险，或发生任何可疑且非预期严重不良事件，须立即报告本委员会
Any amendment requirement which would significantly change the risk of the project, and SUSAR happened in implementation should be report to BGI-IRB immediately
4.跟踪审查日前，须向本委员会提交项目进展报告
Progress report should be submitted to BGI-IRB prior to continuing review
5.结束时须向本委员会提交结题报告
Concluding report should be submitted to BGI-IRB when the project comes to a close

签批 Issued by: 张秀清　副主任委员Vice Chair

张秀清 (signature)

华大基因生命伦理委员会 BGI-IRB (seal)

华大生命伦理委员会
The Institutional Review Board of BGI

2022-6-29

20220418332

批件号 Doc No. BGI-IRB 22069 有效期至 Expired on 2024/04/14

审查文件 Documents Reviewed

★ 审查申请表、诚信及利益冲突声明 Application Form, Integrity Declaration & COI

★ 项目方案(含申请人履历) Project Proposal(with Researcher CV) v4.0 20220621

☐ 知情同意书模板 Informed Consent Form

☐ 推广宣传材料 Advertisement Documents

★ 合作协议 Cooperation Agreement

☐ 合作方伦理意见 Ethical Clearance from Collaborator

☐ 样本来源的支持性文件 Supporting Document for Materials Source/MTA

☐ 结果告知书 Feedback Form

★ 其他 Others

信息库访问修改控制流程

备注 Remarks
